# Land use configuration shapes climate change risk to gallery forests in a savannah ecosystem

**DOI:** 10.1101/2023.09.19.558476

**Authors:** Henrike Schulte to Bühne, Joseph A. Tobias, Sarah M. Durant, Nathalie Pettorelli

**Affiliations:** Institute of Zoology, Zoological Society of London, Regent’s Park, NW1 4RY London, UK; Department of Life Sciences, Imperial College London, Buckhurst Road, SL5 7PY Ascot, UK

**Author notes:** Corresponding author. Telephone: *0044 020 7449 6498*.

**Keywords:** stressor interaction, risk assessment, West Africa

## Abstract

Interactions between anthropogenic pressures make it difficult to predict biodiversity change and plan conservation interventions. Climate change is expected to drive widespread ecological change in the tropics over the coming decades, but it is unclear where and when these changes are going to intensify, or reduce, the impacts of additional pressures from human land use. To address this uncertainty, we apply a novel risk assessment framework to show how land use configuration modifies risks arising from climate change to gallery forests, an important vegetation type in tropical savannahs. Our analysis shows that the spatial distribution of climate change (specifically, change in annual rainfall) interacts with the spatial distribution of land use (specifically, cropland), as well as the biophysical context of the study site (the W-Arly-Pendjari transboundary protected area in West Africa), to shape the risk that changes in rainfall pose to gallery forests in the region. Due to the pathways by which rainfall change and land use interact, risks are especially elevated in core protected areas, warranting particular attention from conservation managers. Overall, our work illustrates how unexpected patterns in risks can arise through interactions between pressures on biodiversity, highlighting the importance of considering mechanistic pathways for predicting biodiversity outcomes under multifacetted global environmental change.

## 1. Introduction

Understanding and predicting ecological responses to multiple anthropogenic pressures is key to managing biodiversity in the 21^st^ century (França et al. 2020, Brown et al. 2013). It is currently challenging to anticipate changes in biodiversity in situations when multiple pressures are present because any given pressure can modify the effects of another on biodiversity, resulting in unexpected biodiversity outcomes. This is a problem for conservation decision making for two reasons: first, pressure interactions can make it difficult to choose between alternative conservation strategies, as their efficacy can vary depending on interaction outcomes (Brown et al. 2013); second, pressure interactions can alter risk rankings of different sites, and thus spatial prioritisation outcomes for conservation intervention (Brown et al. 2014). Climate change and land use change are two particularly pervasive agents of biodiversity change (Newbold et al. 2016, Pecl et al. 2017) which have been shown to interact, resulting in unexpected ecological outcomes (Oliver & Morecroft 2014, Côté et al. 2016). However, predicting the outcomes of climate change in landscapes modified by human land use has been limited so far by a lack of mechanistic understanding of how these interactions shape biodiversity (Schulte to Bühne et al. 2021).

Gallery forests in tropical savannah ecosystems are at a particularly high risk of interactions between climate change and the surrounding land use. Gallery forests are wooded areas along permanent and temporary rivers, as well as topographic depressions that never carry surface water, in tropical savannahs that are otherwise characterised by a more open, or no, tree canopy (Silva et al. 2008, Müller et al. 2012, Azihou et al. 2013). The tree assemblage in gallery forests is distinct from woody species found in the surrounding savannah, with gallery forest species typically more sensitive to fire (Gignoux et al. 2009, Armenteras et al. 2021). Gallery forest species can locally outcompete more fire-resistant competitors in riverbeds and other topographic depressions because these areas have enough water and nutrients to allow gallery forests species to grow fast and shade out savannah species, especially grasses (Hoffmann et al. 2009). The exclusion of highly flammable savannah grasses reduces the incidence of fire, leading to a positive feedback loop between gallery forest species regeneration and fire suppression. Given that climate change will likely alter water availability and fire frequency in the tropics (Ranasinghe et al. 2021), and consequently the competitive ability of the fire-resistant plant communities surrounding gallery forests, gallery forests can be expected to be highly sensitive to predicted changes in climate (Behling et al. 2005, Leal et al. 2016).

In addition, gallery forests are typically long and thin (Azihou et al. 2013), meaning that any edge effects from surrounding human land use can potentially affect the majority of their area, and may thus play a key role in shaping the response of gallery forests to climate change. Gallery forests often occur in areas characterised by agricultural expansion (e.g. Solefack et al. 2018); in addition, gallery forests themselves are often used as sources for fire wood, fruit or other non-timber forest products (Ahouandjinou et al. 2019, Solefack et al. 2018). However, it is currently unclear where such land uses may increase or decrease risks arising from climate change to gallery forests, making it difficult to prioritise areas for conservation intervention. This is a problem because, despite their relatively small extent, gallery forests are crucial for the overall functioning of savannah ecosystems, e.g. providing dry season refuges for birds and amphibians as well as habitat for forest-dwelling species, and storing significant amounts of carbon (Johnson et al. 1999, Natta et al. 2002, Dimobe et al. 2018, Silva et al. 2008).

Generating information about where gallery forests are at a high risk of adverse effects due to climate change-land use interactions is difficult because (1) the effects of climate change-land use interactions are diverse and context-specific (Mantyka-Pringle et al. 2012), and (2) they require a considerable amount of data to quantify at scales appropriate to conservation management (Oliver & Morecroft 2014). To address this knowledge gap, we here adapt for the first time a recently developed risk framework that explicitly includes the ecological effects of interactions between climate change and surrounding land use (Schulte to Bühne et al. 2021) to quantify the risks that climate change-land use interactions pose to gallery forests. This framework starts with creating a context-tailored conceptual model of how land use may modify different dimensions of climate change risks to gallery forests (Figure 1), and then uses open-access data to quantify these interactions. This allows mapping relative risk to this habitat type despite the absence of context-specific quantitative assessments of climate change-land use interactions, thus enabling the rapid generation of decision making-relevant insights into future risks from climate change to gallery forests, and how these are shaped by the spatial distribution of land use.

**Figure 1:**
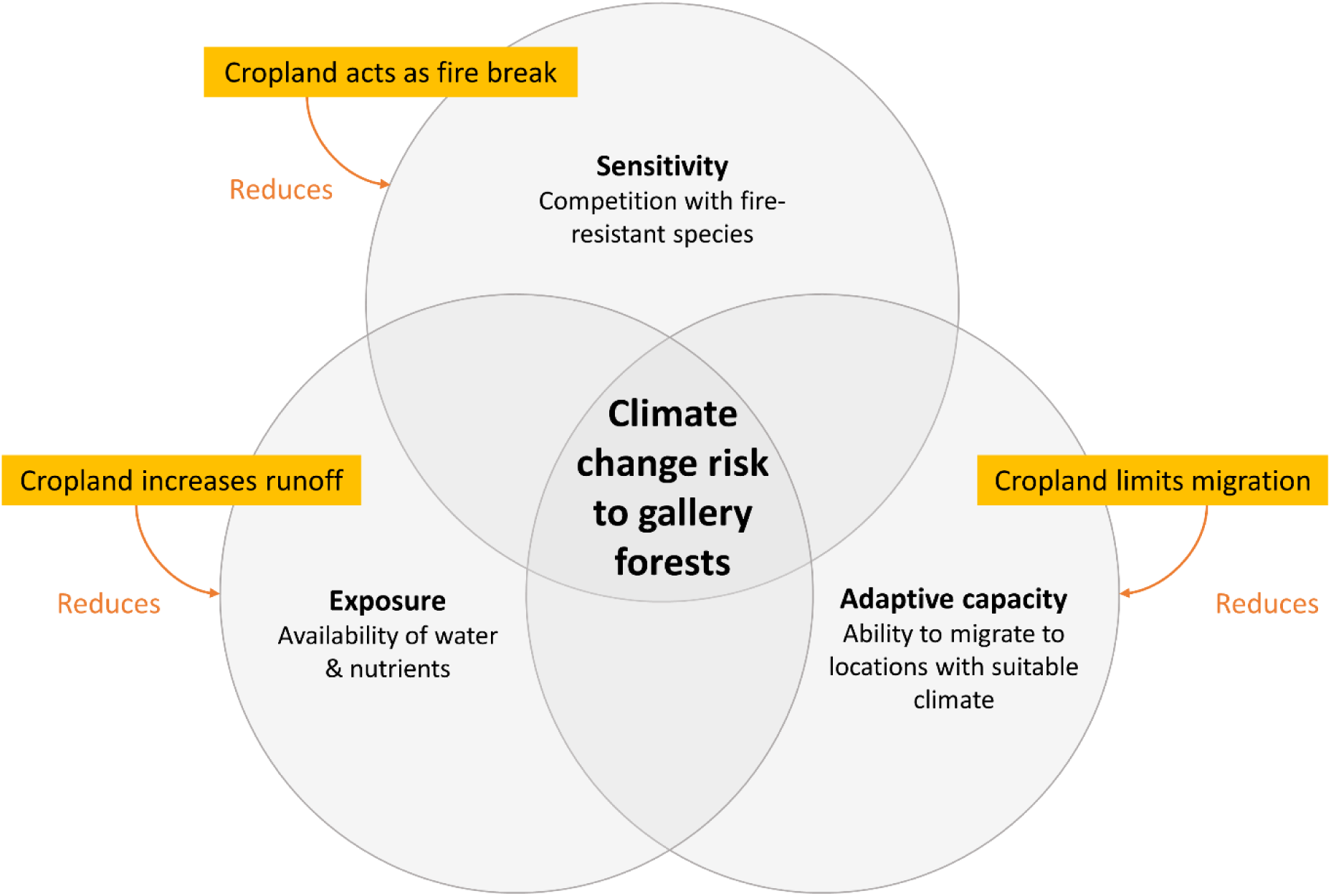
Overview over the mechanisms that shape the risk of climate change to gallery forests, and how each risk dimension (exposure, sensitivity, adaptive capacity) is modified by land use in the form of agriculture.

## 2. Methods

### 2.1 Overview over study site

The W-Arly-Pendjari transboundary protected area complex (WAP), a 35,400 km^2^ network of protected areas in West Africa, is a suitable site to evaluate the risks of climate change-land use interactions for gallery forests (Figure 2). It is the largest contiguous savannah landscape in West Africa (Amahowé et al. 2013) and has in the past experienced significant conversion of savannah vegetation (including gallery forests) to cropland, predominantly outside of protected areas (Schulte to Bühne et al. 2017). This has led to the isolation of habitats across the WAP from other savannah areas in the region (Clerici et al. 2007). The protection of gallery forests is seen as a key management aim in this region (CENAGREF 2014, 2015). Climate change will be posing an additional threat to this biodiversity hotspot over the next century (IPCC 2021), creating the conditions for potentially significant climate change-land use interactions.

**Figure 2:**
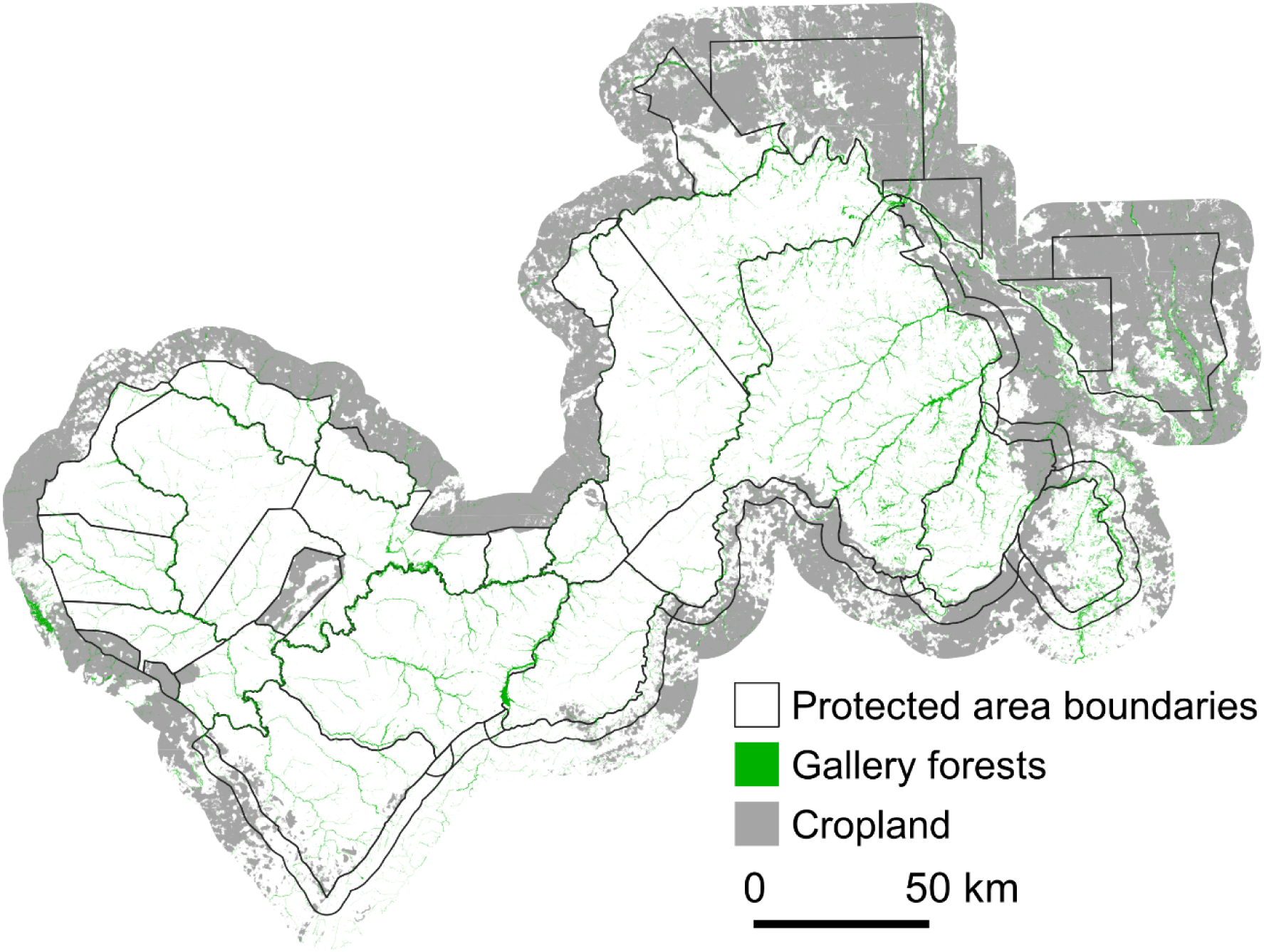
Overview over the study site, the W-Arly-Pendjari transboundary protected area complex, showing the distribution of gallery forests.

### 2.2 Overview over risk assessment framework

First, gallery forests in and around the WAP were mapped based on remote sensing data (Section 2.3.). Then, key mechanisms shaping exposure, sensitivity, and capacity of gallery forests to adapt to future climate change were identified from the scientific literature (Section 2.4, Figure 1). Appropriate indicators were defined for each mechanism, and the overall risk from climate change for existing gallery forest habitat across the WAP was evaluated for two contrasting climate change scenarios (“wet” and “dry”). Then, a conceptual model was created to articulate how the current distribution of agriculture change alters each climate change risk component, again based on existing literature (Section 2.5). Finally, we evaluated how the overall risk from future climate change to gallery forest is affected by current human land use (Section 2.6).

### 2.3 Mapping gallery forests across the study site

Gallery forests were mapped across the study site using a supervised classification approach based on Sentinel 2 data from 2020 and 2021 (for details, see Supplementary materials). The resulting dataset contained 608,756 gallery forest patches. To speed up processing, further analysis was restricted to patches of more than 1,000 m^2^ (i.e. > 10 Sentinel pixels), given that small patches are more likely to be commission errors. The remaining 86,442 patches represented 91% of the original gallery forest area, meaning the majority of the identified gallery forest area was retained for analysis while significantly reducing downstream computational requirements.

### 2.4 Identifying and quantifying climate change risk

Risk is defined as the likelihood of an adverse outcome that negatively affects biodiversity (Reisinger et al. 2020). Here, we focused on the risk of loss of extent of gallery forest in the WAP due to climate change, but it should be noted that loss of species diversity or ecological function of remaining gallery forests would also be classed as adverse outcomes. Risk, in the context of this study, only refers to gallery forests. The loss of savannah species due to invasion by gallery forests could be described as an adverse outcome, too, from the perspective of the savannah species community; however, this is not the focus of this work, given the relatively large extent of savannah compared to gallery forests in our study site.

#### 2.4.1 Exposure

Gallery forests establish where water availability is locally greater than in the surrounding landscape, due to access to groundwater and/or high soil moisture. Both groundwater levels and soil moisture tend to increase with rising rainfall levels (Fontaine et al. 2007, Richard et al. 2013, Cuthbert et al. 2019, 2019b, MacDonald et al. 2021), suggesting that the exposure of gallery forests to climate change depends on the magnitude of changes in rainfall they experience. To quantify exposure to climate change, the relative change in annual rainfall between the present period (2000-2018) and the future (2081-2100) was calculated based on historical monthly precipitation data from WorldClim v2.1 (at 2.5° degrees spatial resolution, Fick & Hijmans 2017) and future rainfall predicted by CMIP6 projections (CRU-TS 4.03, Harris et al. 2014) downscaled and calibrated based on the WorldClim v2.1 data (Fick & Hijmans 2017; Table 1). Rainfall predictions across West Africa are highly uncertain, so to characterise the range of potential climate futures, two contrasting scenarios of future rainfall (BCC-CSM2 and CanESM5, both Shared Socio-economic Pathway 585) were chosen to quantify climate change risk. These scenarios showed the strongest declines and increases, respectively, for rainfall across the study site (called dry and wet scenario, respectively, in the following).

**Table 1:** Overview over data sources used to map gallery forests and quantify risks from climate change and land use.

| Data | Description |
| --- | --- |
| Gallery forests | Land cover classification based on Sentinel 1 and 2 data, groundtruth data from Google Earth |
| Elevation | Shuttle Radar Topography Mission 1 Arc-Second Global dataset from USGS EarthExplorer |
| Topographic depression network | Identified based on the elevation data, using the r.watershed algorithm in QGIS |
| Predicted rainfall change | Difference between predicted and current annual rainfall (WorldClim v2.1 for current rainfall, and CMIP6 projections for future rainfall) |
| Distribution of cropland | Land cover classification based on Landsat 8 data, groundtruth data from Google Earth |

In the dry scenario, all pixels across the study site experience a decline in rainfall. Values were standardised so that a change in rainfall of 0% corresponds to a risk score of 0 (no decline in rainfall), and a score of 1 corresponds to the maximum observed relative change in rainfall (ca. −30%). A higher score thus indicates a higher likelihood of an adverse outcome (loss of gallery forest extent). By contrast, in the wet scenario, all pixels are predicted to experience an increase in rainfall. Values were standardised so gallery forests expected to experience a predicted rainfall change of 0%, which have no opportunity for climate-driven expansion, were assigned a risk score of 1 (denoting the highest risk of an adverse outcome, i.e. no expansion). Conversely, gallery forests predicted to experience the largest rainfall increase (ca. +91%) were given a risk score of 0, indicating that these forests have the largest potential for climate-driven expansion (and thus the lowest risk of an adverse outcome). Each gallery forest patch was assigned an exposure score based on the rainfall change pixel in which the majority of its area fell. Most gallery forest patches are smaller than the resolution of the underlying rainfall data, and so fell within a single pixel.

#### 2.4.2 Sensitivity

The local spatial distribution of gallery forests in a savannah landscape is shaped by competition between slow-growing, fire resistant savannah grasses and woody plants with fast-growing, fire sensitive gallery forest species. Specifically, gallery forest can establish and persist in areas with locally high water and nutrient availability, such as gullies and topographic depressions (Richard et al. 2013), where they are able to grow fast enough to establish a closed canopy. This leads to the suppression of C4 grasses and thus limits the volume of fire fuel (Natta et al. 2002, Azihou et al. 2013), creating a positive feedback loop, where fire-sensitive species inside the gallery forests benefit from reduced fire incidence (and vice versa). Consequently, the response of a gallery forest patch to changes in climate will be stronger if that patch experiences competitive pressure from fire-resistant savannah species across a larger interface.

The sensitivity of a gallery forest patch to climate change was measured in two ways: one based on the size of each patch, and one based on the length of its boundary with savannah vegetation. First, we linearly rescaled gallery forest patch size, so that the largest patch (30.6 km^2^) was assigned a sensitivity score of 0, indicating the lowest sensitivity to declines or increases in rainfall, while the smallest patches (0.001 km^2^) were assigned a sensitivity score of 1, indicating the highest sensitivity. This reflects the assumption that smaller gallery forest patches will be more susceptible to invasion from fire-resistant savannah vegetation as rainfall declines, and conversely that small gallery forest patches are less able to expand into neighbouring savannah vegetation if rainfall increases.

Secondly, the perimeter of each gallery forest was measured, and the difference between the observed perimeter and the smallest possible perimeter for a gallery patch of the same size (the perimeter of a circle with the same surface area) was calculated. Since perimeter increases with patch size, this excess perimeter was divided by the gallery forest surface area. Larger excess perimeters indicate a proportionally longer boundary across which competitive interactions can take place, and thus higher sensitivity to both declines and increases in rainfall. Each indicator was thus standardised to range between 0 (low sensitivity) and 1 (high sensitivity) by a linear scale. Finally, the mean of the two standardised indicators was taken as the final sensitivity score for each gallery forest.

#### 2.4.3 Adaptive capacity

Gallery forests can adapt to changes in climate by migrating to areas where climatic conditions are more suitable, most likely by expanding from existing patches (Wigley et al. 2010, Tng et al. 2012). Tree species that are common in gallery forests in and around the WAP (see Table S1, Supplementary Materials) tend to be dispersed by mammals or birds, as well as wind, with a few species using other modes of seed dispersal, such as water or explosive mechanisms. In theory, this means that seeds can be dispersed relatively far, especially if carried by birds and mammals. In practice, given the habitat preferences of gallery tree species and their sensitivity to fire, it is unlikely that gallery forest trees will establish when surrounded by savannah species, meaning that in the short to medium term (ca. 80 years, the temporal scale of this risk assessment), it is reasonable to assume that expansion of gallery forests will mostly occur from existing patches, rather than through the establishment of new patches. We thus assumed adaptive capacity to be determined by the degree with which existing gallery forests can expand, rather than by the potential for new patches to be established.

Because gallery forests are dependent on locally elevated water availability, expansion from existing patches is constrained by the local topography, and can only occur along topographic depressions. Estimates of expansion rates of gallery forests in response to past climate change vary widely, from ca. 150 m to up to 740 m per year (Watrin et al. 2009), meaning that within 80 years, gallery forests could conceivably shift by between 12 km and 60 km. We arbitrarily chose a distance of 10 km as a conservative estimate of the potential distance that gallery forests could shift over the time period considered here and identified, for each gallery forest, its future position if it migrated in a straight line in the direction of the climate change shift (ideal future position). The direction of the climate change shift was determined by the bearing of an imaginary line between the current position of a gallery forest, and the closest pixel whose future rainfall is identical to the current levels experienced by this patch. “Identical” was defined as having equal annual precipitation when rounded to the first decimal point. We then identified the closest point to this ideal future position in the topographic depression network directly linked to the position of the gallery forest patch (realised future position) and measured the distance between them (Table 1). If the distance between these points was large, this indicated low adaptive capacity, as the topographic network constrains the ability of a given gallery forest patch to migrate in the direction of climate change, in both the wet and the dry climate change scenario. If the distance between them was small, this showed high adaptive capacity.

We also calculated the distance each forest patch would have to migrate along the topographic depression network to arrive at the realised future position. Again, large distances reflect low adaptive capacity (as expansion from a given gallery forest has to overcome a large distance to result in gallery forests being present in an area with suitable climate), and vice versa. Both indicators were standardised, with low values corresponding to 0 (low non-adaptivity, meaning low risk of adverse outcome) and high values corresponding to 1 (high non-adaptivity, meaning high risk of adverse outcome). The final adaptive capacity score was the mean of the two standardised indicator values.

### 2.5 Identifying and quantifying adverse effects of land use on climate change risk

The definition of “adverse effect” of land use depends on the overall effect of climate change on gallery forests. If climate change has an overall negative effect on gallery forests, agricultural land use will exacerbate this risk if it increases the likelihood of loss of gallery forest extent for a given unit of climate change. However, in cases where climate change overall benefits gallery forests (such that increases in extent are expected to occur), agricultural land use will increase climate change risk if it decreases the likelihood of gallery forest expansion for a given unit of climate change. Given that land use change in the form of cropland expansion in this landscape has been, in the past, limited by protected area designations (especially strict designations such as National Parks; Schulte to Bühne et al. 2017), we assumed that future cropland expansions are negligible, and evaluated the interaction between future climate change and the current spatial distribution of land use.

#### 2.5.1 Exposure

Land use in the form of cropland can affect the exposure of gallery forests to changes in rainfall by increasing surface water runoff, which results in higher soil moisture levels and rising groundwater tables, even when overall rainfall is declining (Favreau et al. 2011, Owuor et al. 2016). This means the presence of cropland likely reduces the exposure of nearby, downstream gallery forests to declining rainfalls, as they receive a higher proportion of upstream precipitation. Conversely, it likely increases the exposure of nearby downstream gallery forests to increases in rainfall.

The change in exposure to climate change due to agricultural land use was quantified by the distance of a given gallery patch to the closest cropland patch that was located upstream of the forest. The distribution of cropland was mapped using Landsat 8 satellite imagery from the wet and dry season of 2016/17, using a supervised classification approach that differentiated cropland from non-cropland areas (see Supplementary Materials, Table 1).

In the dry scenario, the exposure of gallery forests to potentially harmful declines in rainfall increases with distance to upstream cropland. Because of this, we used standardised distance values of 1 to characterise largest distances (45 km), and standardised distance values of 0 to characterise gallery patches that are located on cropland (0 km). Similarly, in the wet scenario, exposure to potential beneficial increases in rainfall decreases with distance to upstream cropland, meaning that large distances from upstream corresponded to higher risk of adverse outcome (i.e., no or lower increase in net gain).

#### 2.5.2 Sensitivity

Areas in the WAP that are close to cropland experience some fire protection due to the fuel break effect of cropland (Schulte to Bühne et al. 2023). Such fire suppression likely benefits gallery forests, meaning that gallery forests close to agricultural areas are likely to be less sensitive to adverse climate-mediated changes in fire dynamics. This benefit is expected to decline from a maximum at 0 km to a plateau at ca. 20 km (based on visual inspection of the relationship between fire size and distance to cropland modelled in Schulte to Bühne et al. 2023; see Figure S1 in the Supplementary Materials). Adverse changes in fire dynamics are possible both in the dry and the wet scenario: In the dry scenario, the size of fires that occur later in the season, which are more intense and thus likely more harmful to gallery forests, is likely to increase; conversely, in the wet scenario, the size of early fires is likely to increase (which could also harm gallery forests). Consequently, we assumed the effect of nearby cropland, namely the reduction of climate-mediated changes in fire dynamics, to be beneficial in both climate change scenarios. Changes in gallery forest sensitivity to climate change were quantified as the minimum distance of each gallery forest to the nearest cropland. Because estimated fire size does not increase further for distances from cropland larger than 20 km, the distances of gallery forests that were further than 20 km from the nearest cropland were set to 20 km before rescaling the values to between 0 (low sensitivity, i.e. small distances to cropland) and 1 (high sensitivity, i.e. large distances to cropland).

#### 2.5.3 Adaptive capacity

Assuming that gallery forest expansion will start from existing patches (Wigley et al. 2010), the capacity of gallery forests to adapt to climate change may be curtailed on agricultural land, e.g. due to harvesting of timber or tree fruits (Dimobe et al. 2015, Haarmeyer et al. 2013, M’Woueni et al. 2019). Consequently, if the migration of a given gallery forest patch meets agricultural land, its adaptive capacity is reduced. To quantify this, we measured the distance between the realised future end point of migration (see Section 2.4.3) to the nearest cropland, and compared this to its present distance. For a large number of gallery forests, the difference in the distance was close to 0 (Figure S2, Supplementary Materials), meaning their adaptive capacity is unchanged. We chose 100 m as a conservative cut-off point, accounting for inaccuracies of small distance estimates introduced by the spatial resolution of the cropland and gallery forest maps (30 m). If the distances differed by less than 100 m, we assumed no change in adaptive capacity (giving it a score of 0.5). If the distance increased, we assumed that the adaptive capacity increased (score = 0); conversely, if the distance decreased, the adaptive capacity was assumed to decrease (score = 1). For some gallery forest patches, we were unable to calculate an adaptive capacity score as 10 km migration in the same direction as the isohyets would take them outside the area in which cropland was mapped. For the dry scenario, where isohyets move from North to South, these gallery forests (n = 6145) were predominantly located at the southern border of the study site, and vice versa for the wet scenario (n = 5184). These gallery forests were excluded from the calculations of overall risk scores (see Figure S3 in Supplementary Materials).

### 2.6 Calculating overall risk scores

A climate change risk score was calculated separately for each climate change scenario as the mean of the three separate climate change risk dimension scores (exposure, sensitivity, non-adaptivity). The overall risk from climate change to each gallery forest patch was classified as very low, low, high or very high if it fell in the 25^th^, 50^th^, 75^th^ or 100^th^ quantile, respectively. Similarly, the combined score of land use effects on climate change risk was calculated and classified as very low, low, high or very high (one per climate change scenario). When combining the climate change and land use risk scores, this resulted in a total of 16 combined risk classes. Since the thresholds were arbitrary, the analysis was repeated using stricter thresholds (10^th^, 50^th^, 90^th^ and 100^th^ quantiles respectively).

## 3. Results

Gallery forests at high risk from adverse changes due to climate change (i.e. a risk of loss of extent in the dry scenario, or risk of no gain of extent in the wet scenario), were concentrated in the Southwest of the study area (Figure 3A and B), regardless of the climate change scenario. There were also some high-risk gallery forests in the Northeast, along major rivers and topographic depressions running from the Southwest to the Northeast. Specifically, in the dry climate change scenario, high-risk gallery forests are concentrated along the southern borders of the main rivers, i.e. along lower order rivers that do not extend very far to the south. This means that they cannot migrate towards the (more humid) South if rainfall declines. Conversely, in the wet climate change scenario, high-risk gallery forests are found North of the main rivers, from where there would be no possibility for them to migrate further North following the shifting isohyets as rainfall increases.

**Figure 3:**
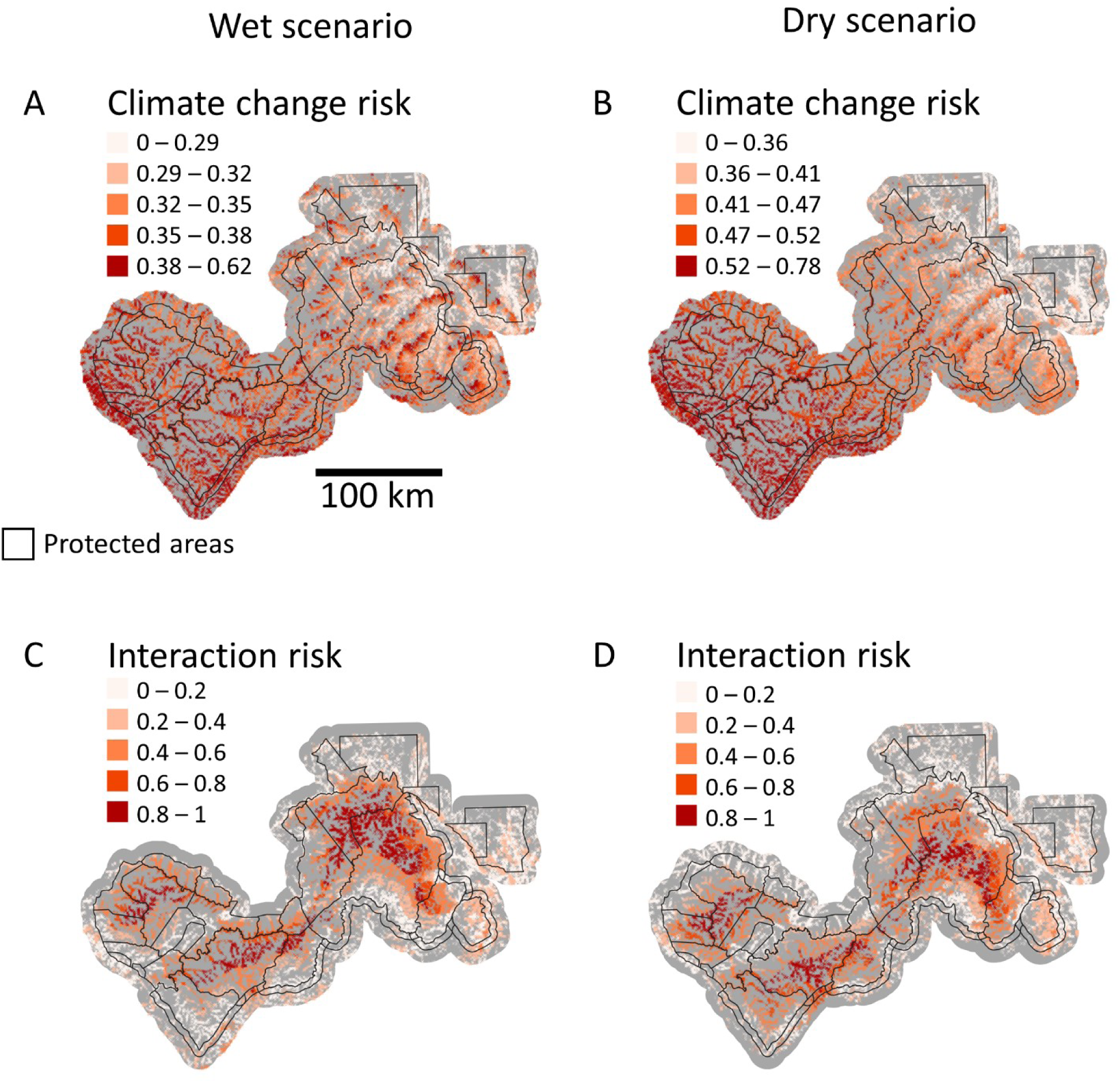
Risks from climate change alone (A and B), and from interactions of climate change with land use (C and D), for the two climate scenarios. Note that for the wet scenario (which assumes increases in rainfall), higher risk scores reflect a lower likelihood that a given gallery forest will be able to expand in response to the improved climatic conditions. For the dry scenario (which assumes declines in rainfall), larger risk scores reflect an increased chance that a gallery forest will contract.

For both the dry and the wet climate scenario, the current spatial distribution of land use in the form of cropland will mainly increase risk from climate change to gallery forests in the core areas of the protected areas (Figure 3C and D). This is because the mechanisms through which the cropland distribution affects gallery forest responses to climate change primarily act to dampen these responses close to cropland. However, it should be noted that gallery forests usually tend to be cleared on cropland, so expansion of cropland should not be considered a strategy to protect gallery forests from adverse climate change effects. In the dry scenario, the high-risk gallery forests are located more to the South compared to the wet scenario. This is likely again the result of the difference in climate change direction, with cropland to the South curtailing migration in the dry climate change scenario, and cropland in the North curtailing migration in the wet scenario.

When combining the two risk levels, three areas of concern emerge: two in the core protected areas of the Southwest, and a smaller, more diffuse area in the Northeast (Figure 4). This reflects both the concentration of gallery forests at high risk from climate change in the Southwest, as well as the dampening effects of nearby cropland on climate change risks. Conversely, gallery forests in the Northeast of the WAP are at the lowest risk from combined climate change and land use, regardless of the climate scenario or risk thresholds. This reflects both the overall lower levels of climate change risks in this part of the WAP, as well as the fact that distances to the nearest cropland tend to be shorter, meaning more gallery forests benefit from the dampening effects of nearby cropland on climate change effects.

**Figure 4:**
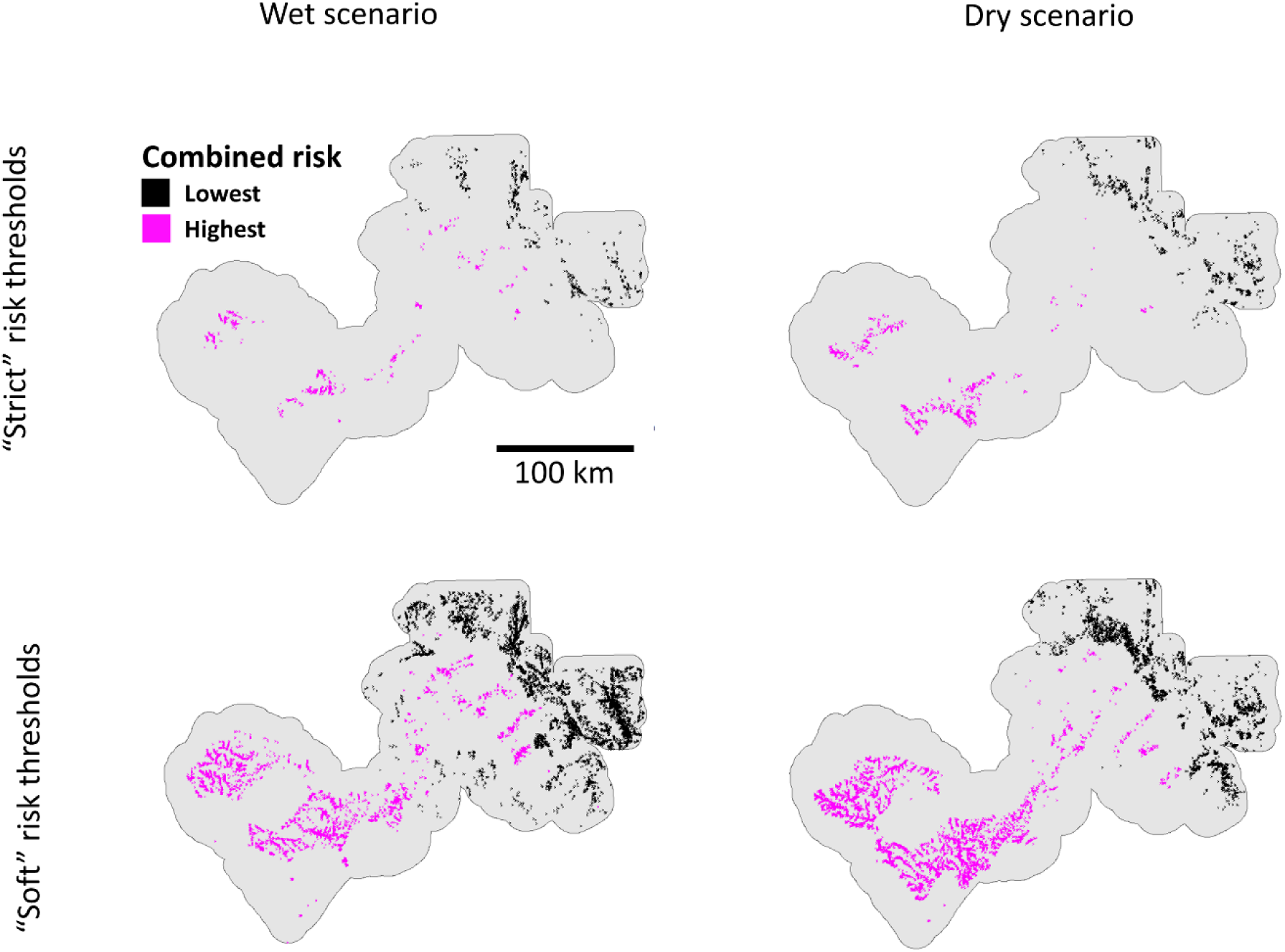
Gallery forests that had both the highest [lowest] climate change and climate change-land use interaction risk index. “Strict” risk thresholds refer to gallery forest that are in the 90^th^ [10^th^] percentile for both climate change and climate change-land use interaction risk. “Soft” risk thresholds refer to gallery forest that are in the 75th^th^ [25^th^] percentile for both climate change and climate change-land use interaction risk. For the wet scenario, which assumes that rainfall will increase, gallery forests with the lowest combined risk are most likely to benefit from the improved climatic conditions and expand, and those with the highest combined risk are least likely to expand. For the dry scenario, which assumes declines in rainfall, gallery forests with the lowest combined risk are the least likely to contract, while those with the highest combined risk levels are the most likely to contract.

Cumulatively, high levels of climate change risk and climate change-land use interaction risk affect comparatively small areas of gallery forests (Figure 5). In particular, the area of gallery forest affected by very high levels of climate change risk that is likely to be exacerbated by the land use context is lower than expected, indicating that the spatial arrangement of land use in the WAP does not systematically exacerbate risks from climate change to gallery forests.

**Figure 5:**
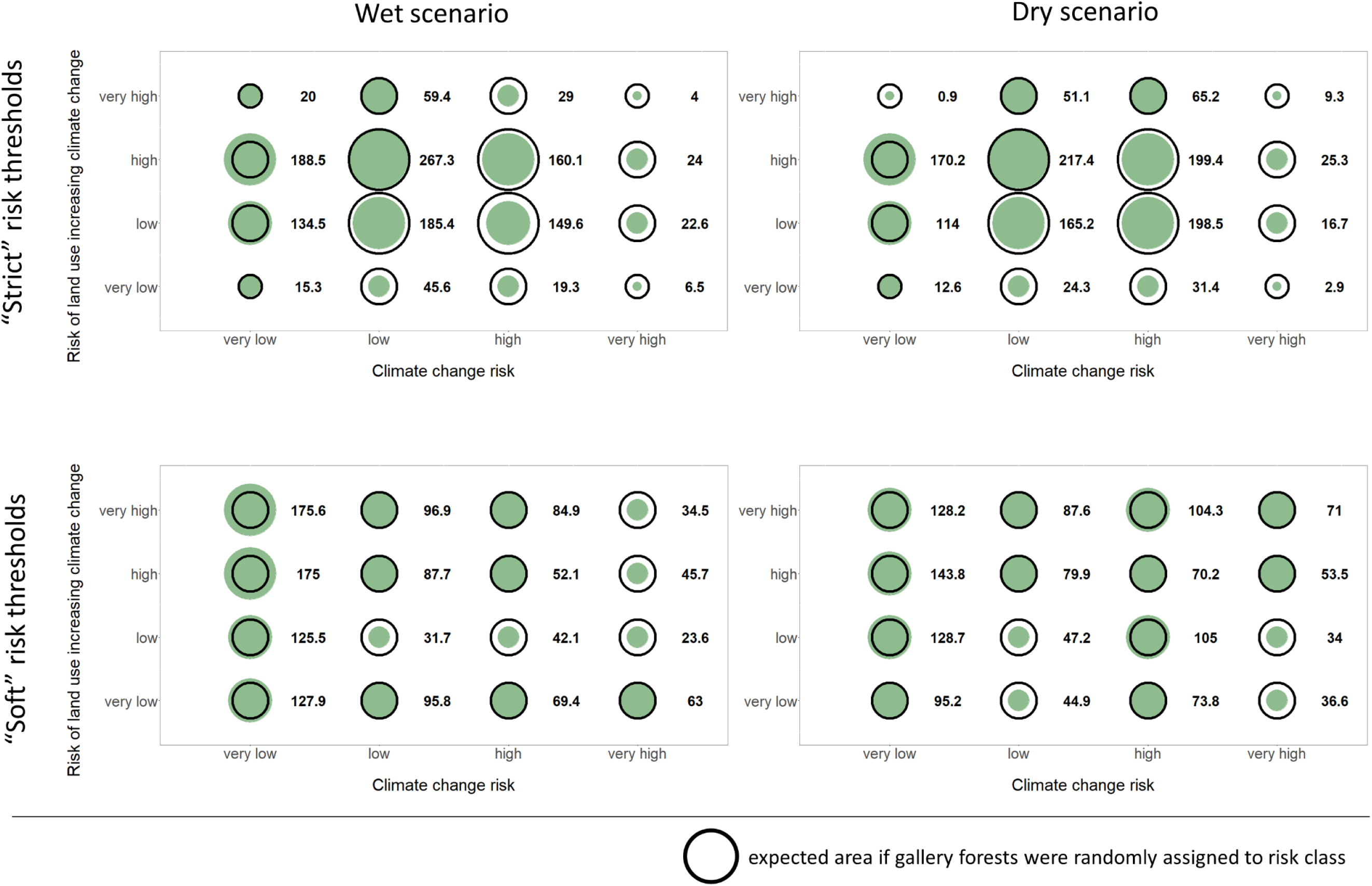
Cumulative area of gallery forest assigned to different risk levels. The numbers refer to the cumulative area of gallery forest in each risk category in km^2^.

## 4. Discussion

In this study, we demonstrate how existing literature and open-source spatial data can be used to assess how the spatial distribution of land use modifies the risks from climate change to gallery forests, an important vegetation type in tropical savannahs. The framework allows (1) identifying gallery forests at high risk of climate change and climate change-land use interactions, and (2) highlighting geographic areas where high-risk (or low-risk) gallery forests cluster. This provides important information to conservation managers and policy makers who need to prioritise stretched resources to mitigate against biodiversity loss in the context of multiple interacting anthropogenic pressures. It also allows contrasting risk outcomes from different climate change scenarios, which are important in situations where climate change projections are uncertain, and where conservation planning will thus have to account for different futures.

If rainfall declines over West Africa, as suggested by the dry scenario, gallery forest conservation will likely focus on preventing or minimising the retreat of gallery forests as water availability declines. Maintaining gallery forests, even if rainfall declines, is possible, as evidenced by the current presence of gallery forests in much drier areas (Moussa et al. 2019). Key to maintaining existing gallery forests will be to minimise the competitive advantage of savannah grasses and trees at the gallery forest-savannah interface (Oliveras & Malhi 2016). Strategies to achieve this include (1) protecting gallery forest from fires, especially late in the dry season, when they tend to be hotter and more destructive (N’dri et al. 2018), e.g. through the application of fire breaks, and (2) protecting gallery forests from significant levels of resource extraction, including browse and fruit (Dimobe et al. 2015, Gaoue et al. 2017) to maintain the performance of existing trees and support regeneration of new cohorts. It will be particularly important to monitor gallery forests in the hotspots highlighted in Figure 4, as these will likely act as sentinels for declines. Conversely, if rainfall increases across West Africa, as suggested by the wet scenario, loss of gallery forest will be unlikely. In this scenario, the risk assessment can help inform the conservation of open areas around gallery forests. Specifically, our analysis suggests that gallery forests are most likely to expand in the North and East of the WAP. In the (unlikely) situation where this may be undesirable from a conservation perspective, fire management could be used to give a competitive advantage to fire-resilient grasses and tree species (Smit et al. 2016).

Our results show that it is the particular arrangement of cropland in the wider landscape (with cropland forming a quasi-ring around the central areas), relative to patterns of predicted climate change, that drives the spatial distribution of risk to gallery forests. Land use patterns are driven by environmental factors such as topography and climate, but also socio-economic, cultural and individual factors, including local agricultural practices (see e.g. Marcos-Martinez et al. 2017, Simmons et al. 2021). Given that these factors can vary widely between locations and over time, it is clear that patterns of climate change-land use interactions will also vary widely, reflecting the often idiosyncratic history of human-environment interactions. This makes it difficult to use pattern analysis alone to improve our understanding of climate change-land use interactions. Instead, a focus on interaction mechanisms allows transferring insights into processes underlying such interactions between study sites (Schulte to Bühne et al. 2021).

Despite the strengths of this risk assessment framework, there are some drawbacks that limit its ability to predict gallery forest change in response to interacting anthropogenic pressures. First, given the limited data and ecological understanding of gallery forests, it was not possible to quantify absolute risk levels (e.g. the absolute probability of a gallery forest disappearing or expanding within a given time unit). To do this, the quantitative responses of gallery forests to different environmental drivers (such as changes in rainfall) would have to be known. However, given that such response rates are often not available (Le Breton et al. 2019), establishing relative risks instead (here: identifying the gallery forests within the study site that are the highest or lowest risk compared to all others) provides important information for conservation of a given management unit, even if these risk levels cannot be directly compared to risks calculated independently for gallery forests at other study sites.

In addition, we limited the scope of the analysis to the risk of a change in extent of gallery forest. This does not account for shifts in gallery forest species composition, or associated changes in functioning, which could have unintended consequences for other species in the wider landscape. Degradation of ecosystem and habitats is often more difficult to detect, conceptually harder to define, and thus more difficult to compare across ecosystems, than changes in extent. This highlights that risk assessments of climate change-land use interactions need to be considered together with multiple other information sources, which may include local and other ecological knowledge, and other qualitative information (McMurdo Hamilton et al. 2021).

Conservation strategies will have to contend with multiple interacting anthropogenic pressures in the 21^st^ century. A climate change-land use change interaction risk framework allows integrating existing knowledge and data about the processes driving such interactions to predict where these interactions are likely going to occur, and where they may require management attention (Saatchi et al. 2021). This framework will contribute to improving the quality and availability of actionable information on complex ecological change, which remains a key challenge of applied ecology.

