## Supplementary Materials for "Land use configuration shapes climate change risk to gallery forests in a savannah ecosystem"

### Supplementary Materials: Supplementary texts

##### Mapping gallery forests across the study site

To identify gallery forests across the study site, all available satellite imagery acquired by Sentinel 2 between 01/11/2020 and 31/03/2021 (corresponding to the dry season) was downloaded (Table 1). After clouds were removed from the imagery, the median Normalized Difference Vegetation Index (NDVI; Pettorelli 2013) was calculated for each pixel (processing was carried out in Google Earth Engine). Sentinel 1 data was also downloaded for the same time period, and the median backscatter in the VV and VH channel calculated for each pixel.

A supervised classification was carried out to identify gallery forest, using the random forest classifier. Groundtruth points (n = 1346) for two land cover types (gallery forests and non-gallery forest) were derived from recent Google Earth imagery and split into training (n = 897) and validation (n = 449) sets. NDVI, VV and VH backscatter, as well as elevation (from Shuttle Radar Topography Mission (SRTM) 1 Arc-Second Global dataset, downloaded from USGS Earth Explorer) were used to train the classifier. The final model had an overall accuracy of 92.4%, with specificity and sensitivity to gallery forests of 92.1% and 93.2% respectively. Processing was carried out in R, using the caret and randomForest packages (Kuhn 2022, Liaw & Wiener 2002).

Contiguous pixels classified as gallery forest were collated into patches. Visual inspection of the gallery forest map revealed that commission errors were due to forested areas that are not gallery forests (i.e. do not grow along a topographic depression) being erroneously classified as gallery forest. To exclude these forested areas, any gallery forest patch that did not overlap with a 120 m buffer along local topographic depression was removed. Local topographic depressions were identified by the r.watershed algorithm implemented in QGIS, based on SRTM elevation data (with depressions removed, Jenson & Domingue 1988).

**Mapping cropland across the study site**

Landsat 8 surface reflectance products from a wet and a dry season (April - October 2016 and December 2016 - February 2017 respectively) were downloaded from the U.S. Geological Survey Earth Explorer. Processing and analysis have been described in Schulte to Bühne et al. (2017). Briefly, tasseled cap-transformed bands and texture metrics were calculated for the dry and wet season data respectively, and added to the original bands for the supervised classification. Groundtruth data was collected from high resolution Google Earth imagery (62 polygons on cropland, 48 on non-cropland). Groundtruth data were converted into spatial points (cropland: n = 294, non-cropland: n = 766). Pixels were classified into cropland and non-cropland using a supervised classification algorithm, with 40% of the training data held back for validation. Again, processing was carried out in R, using the RStoolbox package (Leutner et al. 2019). The overall accuracy of the map was 95.6%, with specificity and sensitivity to cropland at 96.9% and 91.9% respectively.

Visual validation of the cropland map against recent (2020) Sentinel 2 imagery was carried out to delete erroneously classified cropland, especially erroneous patches that were far away from other cropland and would thus introduce large errors in the calculations of distances of gallery forests to cropland. The resulting cropland map reflects the large-scale distribution of cropland across the WAP in the 21$^{st}$ century well (Schulte to Bühne et al. 2017): Most cropland is confined to outside the protected areas, with some important exceptions, such as the Madjoari and Pama enclave (UNDP 2007) and the Zone d’Occupation Controlée along the eastern border of the Pendjari hunting zone (Konrad 2015).

### Supplementary Materials – Tables and Figures

**Table S1**: A list of common tree species in gallery forests in the Sahelian and Sudanian zones in West Africa and their main dispersal modes. The tree species list was compiled from Houehanou et al. 2013, Ganamé et al. 2019, Sambaré et al. 2011, and Kirchmair 2017. “Other” refers to dispersal by insects, by explosive mechanism, and to plants with no obvious primary dispersal mechanism.

| Species | Dispersal mode | Reference |
|---|---|---|
| Khaya senegalensis | wind | Hovestadt et al. 1999 |
| Anogeissus leiocarpa | wind | Hovestadt et al. 1999 |
| Terminalia macroptera | wind | Hovestadt et al. 1999 |
| Mitragyna inermis | wind | Hovestadt et al. 1999 |
| Combretum nigricans | wind | Hovestadt et al. 1999 |
| Pterocarpus santalinoides | wind | Hovestadt et al. 1999 |
| Crossopteryx febrifuga | wind | Hovestadt et al. 1999 |
| Daniellia oliveri | wind | Hovestadt et al. 1999 |
| Combretum micranthum | wind | Hovestadt et al. 1999 |
| Antiaris africana | birds | Hovestadt et al. 1999 |
| Manilkara multinervis | birds | Hovestadt et al. 1999 |
| Syzygium guineense | birds | Hovestadt et al. 1999 |
| Acacia ataxantha | birds | De Swardt & Louw 1994 |
| Acacia seyal | birds | Argaw et al. 1999 |
| Ziziphus mucronata | birds | Zietsman et al. 1989 |
| Piliostigma thonninguii | mammals | Hovestadt et al. 1999 |
| Tamarindus indica | mammals | Hovestadt et al. 1999 |
| Diospyros mespiliformis | mammals | Hovestadt et al. 1999 |
| Chlorophora excelsa | mammals | Hovestadt et al. 1999 |
| Dialium guineense | mammals | Hovestadt et al. 1999 |
| Cola laurifolia | mammals | Hovestadt et al. 1999 |
| Cola cordifolia | mammals | Hovestadt et al. 1999 |
| Carapa procera | mammals, water | Lankoandé et al. 2021 |
| Kigelia africana | mammals | Hovestadt et al. 1999 |
| Elaeis guineensis | animals, humans, water | Hayati et al. 2004 |
| Borasses aethiopium | mammals, other | Bayton 2007 |
| Morelia senegalensis | others | Hovestadt et al. 1999 |
| Berlinia grandiflora | others | Hovestadt et al. 1999 |
| Terminalia laxiflora | not known | |
| Pericopsis laxiflora | not known | |
| Vitex chrysocarpa | not known | |

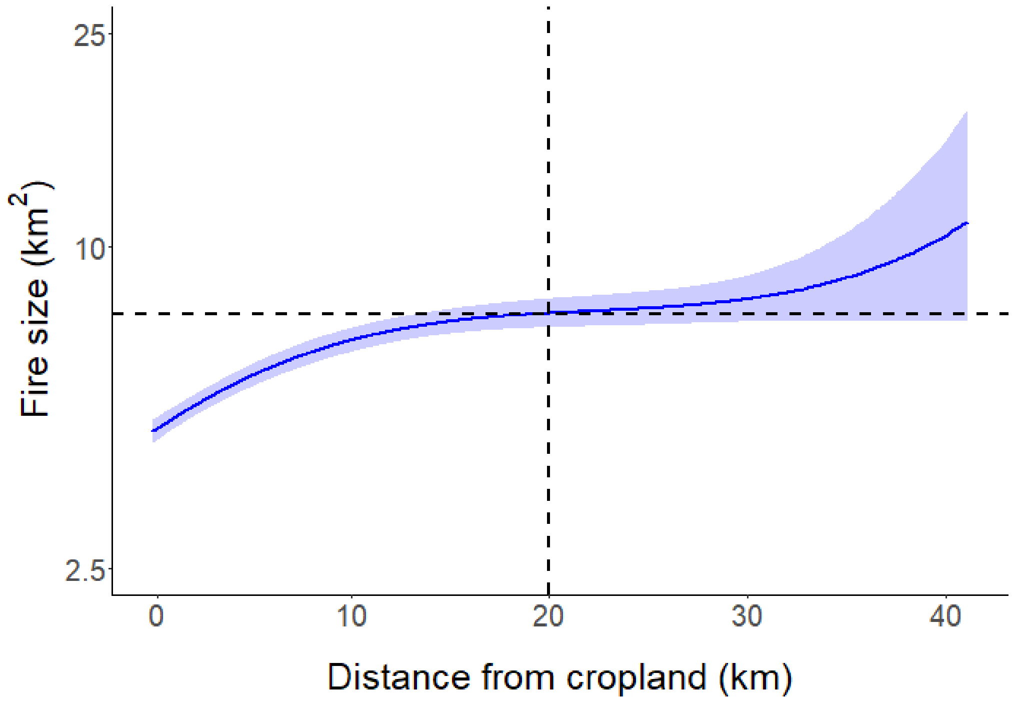


**Figure S1**: The relationship between fire size and distance of fire to the nearest cropland (see Schulte to Bühne et al. 2023 for details of how this relationship was modelled; the shaded area represents the 95% confidence interval). Until a distance of about 20 km, fire size increases as the distance to cropland increases. Then this relationship plateaus, indicating that the fuel break effect of cropland becomes weaker. Fire size rises again at very large distances; however, the confidence intervals also become wider at this point due to a lower number of fires at large distances, so a threshold of 20 km was chosen as a best estimate for the upper limit of the spatial scale of the fuel break effect of nearby cropland.

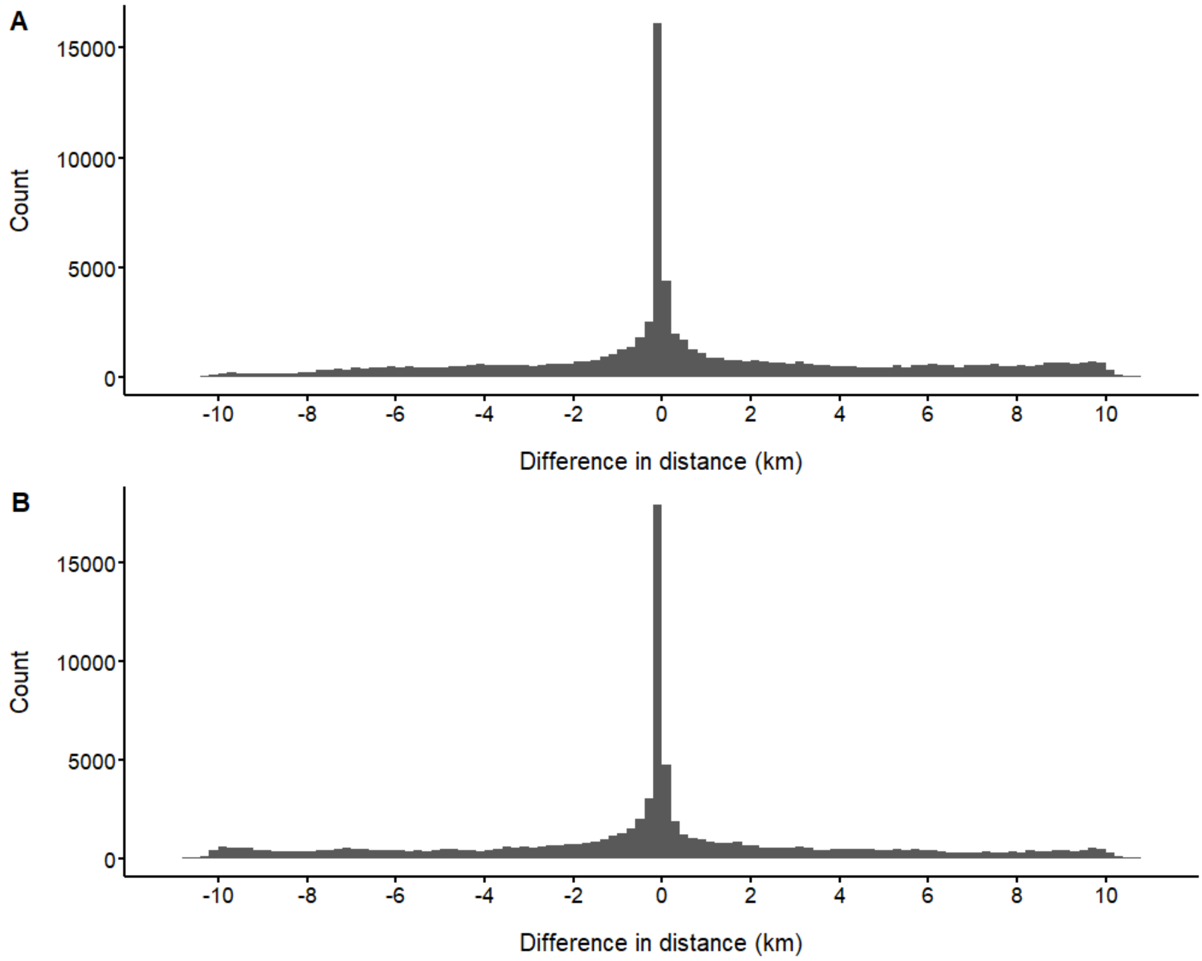


**Figure S2**: Distribution of the difference in distance between each gallery forest and the nearest cropland patch before and after migration of a gallery forest patch in (A) the dry scenario and (B) the wet scenario.

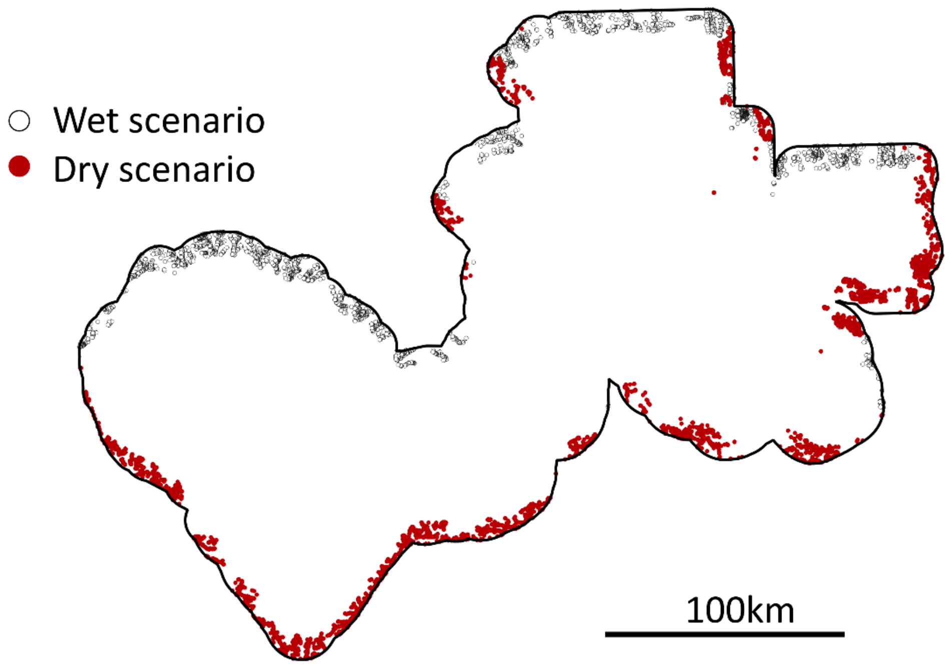


**Figure S3**: Location of gallery forest patches for which it was not possible to quantify the effect of land use on the ability of these gallery forest patches to adapt to climate change by shifting in the same direction as the isohyets, for (A) the dry scenario and (B) the wet scenario. This was due to the boundaries of the cropland map available for this study.
